# A Possible Role Of Microglia In Zika Virus Infection Of The Fetal Human Brain

**DOI:** 10.1101/142497

**Authors:** Julien Muffat, Yun Li, Attya Omer, Ann Durbin, Irene Bosch, Grisilda Bakiasi, Edward Richards, Aaron Meyer, Lee Gehrke, Rudolf Jaenisch

**Affiliations:** Whitehead Institute for Biomedical Research, 9 Cambridge Center, Cambridge, MA 02142, USA; Department of Biology, Massachusetts Institute of Technology, 31 Ames Street, Cambridge, MA 02139, USA; IMES, Massachusetts Institute of Technology, Cambridge MA 02139, USA; Department of Microbiology and Immunobiology, Harvard Medical School, Boston 02115, USA; Harvard-MIT Program in Health Sciences and Technology, Cambridge, MA 02139, USA; Bryn Mawr College, Bryn Mawr, PA; Koch Institute for Integrative Cancer Research at MIT, Cambridge, MA 02139, USA

## Abstract

Maternal Zika virus (ZIKV) infection during pregnancy is increasingly recognized as the cause of an epidemic of microcephaly and other neurological anomalies in human fetuses. However, it remains unclear how ZIKV gains access to the highly vulnerable population of neural progenitors of the fetal central nervous system (CNS), and which cell types of the CNS may serve as viral reservoirs. To model viral interaction with cells of the fetal CNS in *vitro*, we investigated the tropism of ZIKV for different iPS-derived human cells, with a particular focus on microglia-like cells derived from human pluripotent stem cells. We show that ZIKV infected isogenic neural progenitors, astrocytes and microglia-like cells, but was only cytotoxic to neural progenitors. Infected glial cells propagated the virus and maintained viral load over time, leading to viral spread to susceptible cells. ZIKV-infected microglia, when co-cultured with pre-established neural spheroids, invaded the tissue and initiated neural infection. Since microglia derive from primitive macrophages originating in anatomical proximity to the maternal vasculature of the placenta, we propose that they may act in *vivo* as a viral reservoir for ZIKV and, owing to their natural ability to traverse the embryo, can establish infection of the fetal brain. Infection of immature neural stem cells by invading microglia may occur in the early stages of pregnancy, before vascular circulation is established. Our data are also consistent with the virus affecting the integrity of the blood-brain barrier (BBB), which may allow infection of the brain at later stages.

## INTRODUCTION

ZIKV emerged in 1947, and had not been considered a grave threat to humans until very recently (Li et al., 2016a). In 2013, an epidemic in French Polynesia was associated with significant morbidity in the form of Guillain-Barre Syndrome (Cao-Lormeau et al., 2016). Since 2014, the virus has been spreading throughout the Americas and the Caribbean, threatening the southern United States where populations of its mosquito vectors, *Aedes aegyptii* and *Aedes albopictus*, are endemic. Concurrent to the spread of ZIKV in South America, a rise in cases of severe fetal malformations, including microcephaly (Martines et al., 2016a, Mlakar et al., 2016), was rapidly linked to maternal infection, with apparent peak vulnerability during the first trimester of pregnancy (Martines et al., 2016a, Roberts and Frosch, 2016). While neural progenitors have been shown to be highly sensitive to infection, the current epidemic raises many questions. Why is ZIKV presenting so severely and only now? Sequence variations (Wang et al., 2016) between the original 1947 Ugandan strain (ZIKV^u^), and the current circulating variants appear minor. How does ZIKV accomplish its vertical transmission from mother to fetus despite all defenses in place? The damage to the fetus can be quite variable, possibly pointing to narrow windows of opportunity for the virus to wreak havoc on development. What makes human hosts so vulnerable, when mice do not develop the disease unless their antiviral defenses are knocked down (Rossi and Vasilakis, 2016, Mysorekar and Diamond, 2016, Lazear et al., 2016)? This latter consideration questions our ability to model the pathology in rodents, as the needs to dampen interferon responses make the study of viral spread within the mouse host arduous at best. What of the vulnerability of more mature nervous systems: can the virus be completely cleared postnatally, or does it leave neurological sequalae even in absence of overt teratogenicity? Are there circumstances in which ZIKV can reach an adult nervous system from the periphery (Cao-Lormeau et al., 2016, Brasil et al., 2016, Soares et al., 2016)? The consequence of such an event are largely unknown.

In this report, we address some of these questions using *in vitro* modeling to study the interaction of ZIKV with various cells of the CNS. We focused our study on the prototypical ZIKV^u^ strain, which is well characterized and is particularly cytotoxic *in vitro* (Xu et al., 2016). We contrast the tropism and cytotoxicity of ZIKV with that of Dengue (DENV), a closely related flavivirus (Olagnier et al., 2016), to identify key biological differences and steer therapeutic discovery. The interactions between DENV and ZIKV may be particularly relevant, as many patients contracting ZIKV have already been exposed to DENV.

## RESULTS

We generated multiple cell types from human induced pluripotent stem (iPS) cells, allowing derivation from a given donor (Soldner and Jaenisch, 2012, Muffat et al., 2016a), avoiding confounding effects of genetic background on viral infectivity or cytotoxicity. Microglia, neural progenitors, astrocytes and neurons, the four CNS cell types used in our study, were derived from the same pluripotent cells (iPSCs) and grown in serum-free medium (NGD). The cells were infected with the MR766 strain of Ugandan ZIKV (ZIKV^u^), which has been reported to be cytotoxic to human neural progenitors (Tang et al., 2016, Xu et al., 2016). We confirmed that a ZIKV^u^ inoculum corresponding to a multiplicity of infection (MOI) of 1 (as measured independently on Vero cells), was highly infective and cytotoxic to iPS-derived neural progenitors (Fig. 1a, left). The cells accumulated large envelope-positive cytoplasmic inclusions, and died by caspase-mediated apoptosis (not shown). When challenged with increasing multiplicities of ZIKV^u^, NPCs accumulated viral genome (vRNA) up to twice the levels of endogenous *GAPDH* (Fig. 1b, left). When infecting astrocytes, differentiated from NPCs, cells became positive for the ZIKV envelope protein (Fig. 1a, middle), and produced vRNA in excess of endogenous *GAPDH*, but virus-induced cell death or caspase activation were not observed (Fig. 1b, middle). iPS-derived microglia infected with ZIKV^u^ became positive for the ZIKV envelope protein (Fig. 1a, right) and accumulated vRNA in a dose-dependent manner reaching levels of almost 50% of *GAPDH* expression (Fig. 1b, right). To study how neural and glial cells compare to peripheral cells, we exposed skin fibroblasts, endothelial cells and peripheral blood monocytes to ZIKV^u^. As shown in Fig. 1c (left), all cells produced high levels of vRNA, with skin fibroblasts capable of accumulating the highest levels of viral genome (over 10-fold higher than *GAPDH)*, without displaying signs of cytotoxicity.

**Figure 1:**
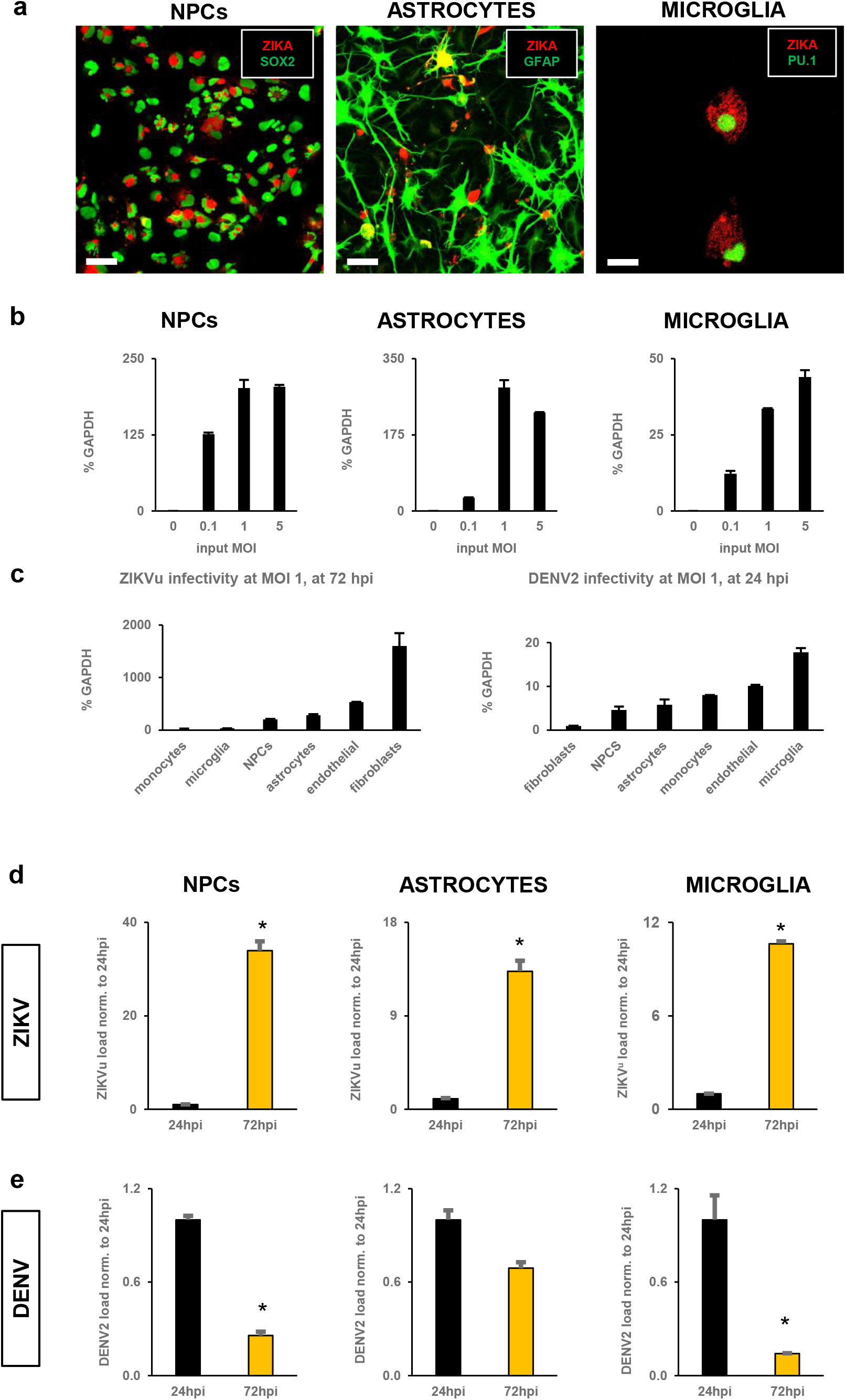
ZIKV and DENV display different tropism and kinetics. **a.** Left panel, Sox2+ (green, nuclear) iPS-derived NPCs infected with ZIKV (red, envelope staining); scale bar 40μm. Middle panel, GFAP+ (green, cytoplasmic) iPS-derived astrocytes derived from the same parental line display cytoplasmic ZIKV inclusions (red, envelope staining); scale bar 40μm. Right Panel, PU.1+ microglia (green, nuclear) derived from the same parental line, also got infected with ZIKV and display numerous cytoplasmic foci positive for ZIKV envelope (red channel); scale bar 12 μm. **b**. dose-dependent infection of different cell types with ZIKV. qPCR of ZIKV RNA (percentage of *GAPDH)* as a function of input inoculum MOI (Left, NPCs; middle, astrocytes; right, microglia). **c**. Left panel, qPCR of ZIKV RNA at 72 hours after infection of indicated cell types with inoculum (MOI 1), as a percentage of *GAPDH.* Right panel, qPCR of DENV RNA at 72 hours after infection of indicated cell types with viral inoculum (MOI 1), as a percentage of *GAPDH.* **d**. kinetics of viral RNA accumulation between 24 hours and 72 hours post infection with ZIKV inoculum (MOI 1) in isogenic NPCs (left panel), astrocytes (middle panel), microglia (right panel). **e**. kinetics of viral restriction between 24 hours and 72 hours post infection with DENV inoculum (MOI 1) in isogenic NPCs (left panel), astrocytes (middle panel) and microglia (right panel). (**b-e**) Values are mean ± s.e.m. (N=2 technical replicates). Asterisks represent significant differences with control, in post-hoc tests after ANOVA (P<0.05).

Dengue (DENV), a flavivirus closely related to ZIKV, is transmitted by the same mosquito vectors but is not known to be overtly teratogenic, though there are indications that it can be neurotropic and neurovirulent (Screaton et al., 2015). To define the different pathogenicity of these related viruses we exposed the same cell types to DENV2. Fig. 1c (right panel) shows that, while the virus infected and replicated in all cell types, the maximum vRNA levels were significantly lower than that of ZIKV, reaching 17% of *GAPDH* levels at their highest in microglial cells. To assess the kinetics of virus replication we compared ZIKV^u^ and DENV2 vRNA levels at different times after infection. As shown in Fig. 1d, ZIKV^u^ vRNA levels increased rapidly from 24 to 72 hours after infection, producing 30-fold more in NPCs (left), 13-fold more in astrocytes (middle), and 9-fold more in microglia (right). In contrast, DENV2 vRNA levels were significantly lower at 24 hours and decreased considerably in astrocytes and microglia over the next two days (Fig. 1e). These results suggest that these cell types limit infection and replication of DENV2, while ZIKV^u^ replication is exponential during this time.

The results in Figs. 1d, e suggest that antiviral responses to infection are particular to either virus, limiting or allowing virus propagation. We assessed the effect of interferon pathways stimulation and inhibition on viral accumulation. When B18R, an inhibitor of type I interferon sensing and signaling, was added to infected cultures, ZIKV^u^ replication was significantly increased after infection of astrocytes and microglia, and only marginally in NPCs (Fig. 2a, yellow bar). Conversely, when infected cultures were exposed to recombinant interferon-γ as an agonist of the type II interferon pathway, virus production was inhibited in all three cell types (Fig. 2a, right bar). In striking contrast, DENV2 did not increase in response to B18R, and was not inhibited by IFN-γ in any of the cell types (Fig. 2b). As shown in Fig. 2c, IFN-γ treatment of ZIKV^u^ -infected NPCs efficiently protected the cells from infection and cell death, raising the possibility that interferon manipulation may be a target of choice to limit cytotoxicity in the fetal brain.

**Figure 2:**
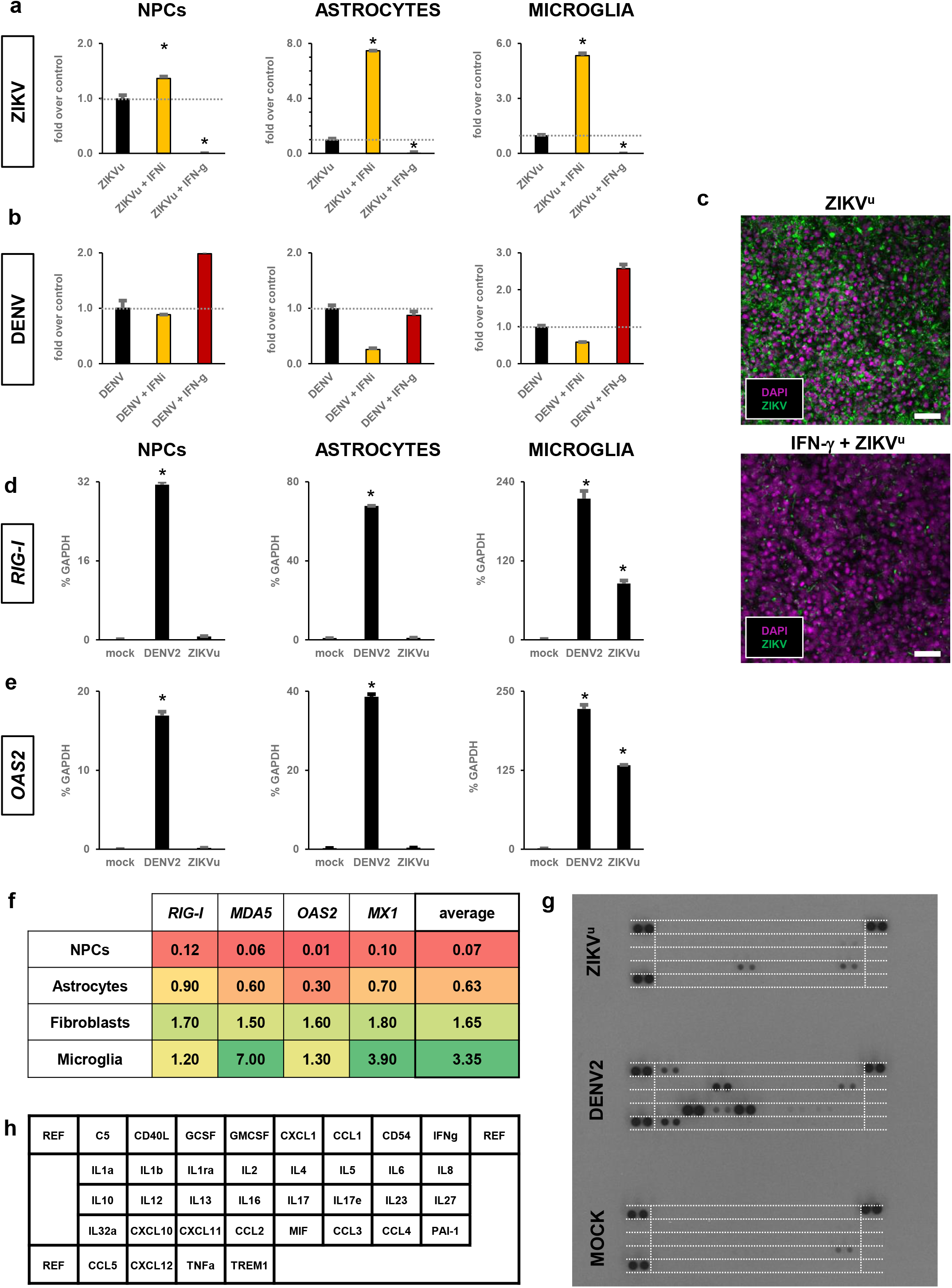
Different interferon responses to ZIKV and DENV infection. **a.** 72 hours after infection, type I interferon inhibition by B18R increased ZIKV load (middle yellow bars), while type II interferon activation by IFN-γ (right bars) decreased ZIKV load (normalized to control ZIKV infection, left black bars and dashed line at 1), in isogenic NPCs (left panel), astrocytes (middle panel), and microglia (right panel). **b**. Effect of type I interferon inhibition (B18R, middle yellow bars) and type II interferon activation (IFN-γ, right red bars) on DENV viral load (normalized to control DENV infection, left black bars and dashed line at 1) in isogenic NPCs (left panel), astrocytes (middle panel), and microglia (right panel). **c**. NPCs infected with ZIKV (top panel, green channel) were protected by pre-treatment with IFN-γ, assessed by reduction in ZIKV signal (green channel), and increase in cell number (nuclear dapi, magenta); scale bars 40μm. **d,e** qPCR for the viral sensor *RIG-I* (**d**) and effector *OAS2* (**e**) following infection with DENV and ZIKV in isogenic NPCs (left panel), astrocytes (middle panel), and microglia (right panel). *RIG-I* and *OAS2* are significantly induced by DENV in all cells, and by ZIKV only in microglia (innate immune cells). Gene expression is expressed as percentage of *GAPDH.* Asterisks represent significant difference with control, in post-hoc tests following ANOVA (P<0.05). **f**. Expression levels of a panel of 4 main effectors of interferon signaling *(RIG-I, MDA5, OAS2, MX1)* and their averages in 4 relevant cell types (NPCs, astrocytes, microglia and skin fibroblasts). Color ranges from red (0% *GAPDH*) to green (7% *GAPDH*:) cells that are hypersensitive to ZIKV (NPCs) have very low basal expression of all 4 genes (average 0.07), while cells that can mount a response to ZIKV (microglia) have high basal expression (average 3.35). **g**. Cytokine profiles of microglia infected with ZIKV and DENV (medium conditioned from 12 hpi to 24 hpi). DENV induced the release of multiple cytokines/chemokines typical of microglial viral activation. In contrast, ZIKV elicited the release of very few cytokines/chemokines (CCL2, IL8), evading innate activation. **h**. map of cytokines and chemokines array probed in **g**, equivalent to the dashed white lines. (**a-b;d-e**). Values are mean ± s.e.m. (N=2 technical replicates). Asterisks represent significant difference with control, in post-hoc tests following ANOVA (P<0.05).

To investigate whether the two viruses elicit different antiviral responses in infected cells we measured gene expression of canonical viral sensors and downstream interferon-stimulated genes (ISGs). RIG-I is a dsRNA helicase acting as a primary sensor of invasion by multiple viruses (Kell and Gale, 2015), including DENV and other flaviviruses. OAS2 is an IFN-inducible enzyme which, in the presence of dsRNA, synthesizes an activator of RNAse L. OAS2 is under interferon control and acts as an effector of viral genome degradation and replication inhibition(Lin et al., 2009). Fig. 2d shows that *RIG-I* was highly induced within 24 hours of exposure to DENV2 (middle bar in each graph) in NPCs (left), astrocytes (middle), and microglia (right) but *RIG-I* levels remained very low in both astrocytes and neural progenitors when exposed to ZIKV^u^ (rightmost bar in each graph) at an MOI that resulted in robust viral infection and replication and caused death of NPCs. This suggests that ZIKV^u^ can escape detection by the canonical antiviral cellular response rendering cells like infected NPCs susceptible to apoptosis. Interestingly, ZIKV^u^ triggered *RIG-I* upregulation in microglia, likely owing to the specialized role of these cells as innate immune effectors. Similarly, *OAS2* was induced in all three cell types by exposure to DENV2 (fig. 2e) and in microglia only by ZIKV^u^.

To assess more broadly the baseline antiviral status of the different cell types, we measured expression of *RIG*-I, *MDA5, OAS2* and *MX1* in absence of infection. As shown in Fig. 2f NPCs, and to lesser extent astrocytes, share low levels of expression of these antiviral response genes, unlike fibroblasts and microglia. Fig. 2g shows that microglia, exposed to DENV2 (middle), expressed and secreted cytokines and chemokines whose combined action could be further involved in neuro-inflammatory processes (such as CXCL10, see blot map in 2h), and elicit cytotoxic effects, in an attempt to defend the host.

To understand virus pathogenesis, it is crucial to define the cell type that initiates and propagates virus in the fetus after primary infection of the mother. As a surrogate for virus production, we tested the supernatant of infected microglia, astrocytes and NPCs for the presence of vRNA. As shown in Fig. 3a, the supernatants of microglia, astrocyte and NPCs infected with either DENV2 or ZIKV^u^ contained vRNA after 72 hours, consistent with virus shedding and with ZIKV^u^-infected NPCs producing their highest levels before death. We observed that DENV2 was cytotoxic to microglia, in contrast to ZIKV^u^, which did not induce death of microglia within 3 days of infection (Fig 3b) but rather resulted in release of viral genomes more than a week after initial infection (Fig. 3c). Large concentrations of ZIKV^u^ vRNA could be detected more than a week after initial infection of astrocytes, while DENV2 was significantly lower (Fig. 3d).

**Figure 3:**
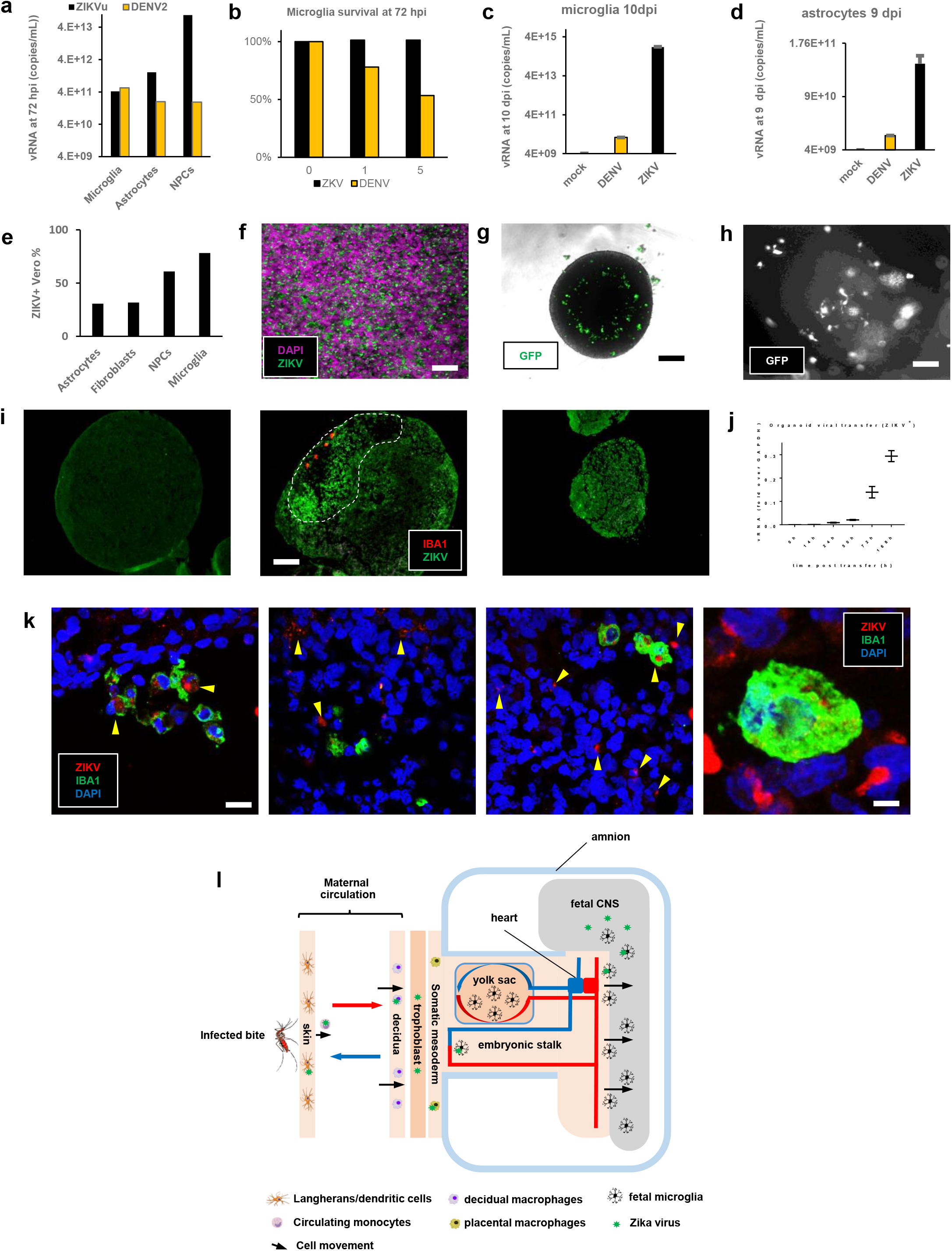
Glial vulnerabilities delineate viral paths to the nervous system. **a.** Viral RNA was present in the supernatant of microglia, astrocytes and NPCs 72 hours post infection. **b**. ZIKV-infected microglia survive infection, while the viability of DENV infected microglia was compromised after 72 hours, in a dose-dependent manner (1 technical replicate, experiment repeated at least twice). **c,d**. Viral RNA was still present in the conditioned medium more than a week after infection of microglia (**c**, 10 days) and astrocytes (**d**, 9 days). ZIKV infected cells, particularly microglia (4 logs), maintained higher levels than cells infected with DENV. Values are mean ± s.e.m. (N=2 technical replicates). **e**. Infectious viral particles were actively produced, as measured by re-infection of interferon-deficient Vero cells by conditioned medium from indicated cell types (N=1). **f**. supernatants from infected microglia efficiently transferred ZIKV (green channel) to a lawn of neural progenitors (magenta, nuclear dapi); scale bar 80μm. **g**. Uninfected microglia (GFP-labeled) co-cultured in suspension with cerebral organoids (spheroids 14 days after reaggregation of NPCs) readily invaded deep in the tissue, mimicking neural tube invasion by microglial precursors; scale bar 200μm. **h**. Uninfected resident microglia adopt ramified morphologies typical of more mature cells; scale bars 20μm. **i**. Section of uninfected cerebral organoid showing background green fluorescence in absence of virus (left panel). When co-cultured with ZIKV-infected microglia (red channel), ZIKV positive signal spreads through the organoid (dashed area), particularly surrounding the invading cells (middle panel). For comparison, an organoid directly exposed to a high titer inoculum becomes much smaller and highly positive for ZIKV (green, right panel); scale bar 100μm. **j**. qPCR of ZIKV^u^ viral amounts as fold over *GAPDH*, in neural organoids paired with infected microglia at t=0h. ZIKV is amplified exponentially in the target organoid. Whisker plots represent mean ± s.e.m. (N=2 technical replicates). **k**. Left panel: confocal pictures of organoid sections showing initial penetration of ZIKV infected (red channel) microglia (Iba1, green channel), into the organoid tissue. Middle two panels: ZIKV viral factories (red channel, yellow arrowheads) were found in the cytoplasm of many cells deep within the organoids. Note: these cells are not the invading cells (green microglia); scale bar 12μm. Right panel: high magnification confocal imaging of organoid host cells directly surrounding an invading microglial cell, clearly showed the presence of ZIKV positive factories in the neural parenchyma of the organoid; scale bar 4 μm. **l**. Diagram of the possible maternal-fetal myeloid chain of ZIKV transmission. Our results indicate that microglia have the unique potential of completing a myeloid chain of transmission, from skin macrophages all the way to the fetus. Pools of infected decidual maternal macrophages wear down placental defenses. Placental fetal resident macrophages get infected but do not travel or come in physical contact with the fetal CNS. Microglial precursors, migrating past the placental and vitelline exchange endothelia, get infected on their way to the embryo, invading the neural tube at its most vulnerable stage. Migration eventually stops, and the blood-brain barrier closes. ZIKV infected immune cells may constitute a viral reservoir for prolonged periods. In contrast, DENV may never reach the fetal neighborhoods, actively restricted at every step in the maternal host, or triggering cell death of microglia.

To test for the presence of active virus, we exposed interferon-deficient Vero cells to the supernatant of the infected human cells. Fig. 3e shows that the supernatant from ZIKV^u^-infected microglia was efficient in establishing infection of Vero cells. When neural progenitors, thought to represent the most relevant target cells for teratogenesis, were exposed to the supernatants of infected microglia, they became homogenously positive for ZIKV envelope protein (Fig. 3f) consistent with the possibility that microglia could initiate infection of the fetal brain.

To model the ability of infected microglial precursors to migrate into the developing CNS and spread ZIKV to vulnerable cells, we developed a 3D-culture model expanding on our initial work replicating an organotypic environment for microglia (Muffat et al., 2016a, Muffat et al., 2016b, Li et al., 2016b). Fig. 3g, shows that immature GFP-labeled microglia, paired with a neuralized organoid, actively migrate into the growing tissue, taking residence and adopting the ramified morphology of more mature microglial cells (Fig. 3h). When microglia that were pre-infected with ZIKV^u^ were co-cultured with these organoids (Fig. 3i, middle panel), we found that invading microglial cells adopted a morphology reminiscent of Gitter cells within 5 days, as large amoeboid cells filled with granular phagocytic content (Das, 1976). Surrounding the invading microglia (red channel), we observed a zone of loose cellular structure displaying elevated ZIKV envelope signal interspersed with pyknotic and fragmented nuclei, indicative of an ongoing degenerative process (green, dashed loop). For reference, we showed that direct infection with a ZIKV inoculum of a similar-sized organoid, resulted in stalled growth, as previously reported (Garcez et al., 2016, Qian et al., 2016) (Fig. 3i, right panel). Fig. 3j depicts the increase in viral load in the target organoid, in the days following pairing with infected microglia. Higher magnification of organoid sections showed that the ZIKV-positive microglial cells initiated infection of adjacent cells, as exemplified by the large ZIKV-positive cytoplasmic inclusions found in nearby parenchymal cells (Fig. 3k, right panel, yellow arrowheads, all panels).

Astrocytes are, in intimate partnership with endothelial cells, the primary components of the blood-brain-barrier (BBB) defining the specialized immune environment of the CNS (Obermeier et al., 2016), preventing entry of pathogens and immune cells. We compared viral cytotoxicity to astrocytes and endothelial cells, exposing homogenous cultures to ZIKV^u^ and DENV2. As shown in Fig. 4a (middle panel) DENV2 exposure was cytotoxic to astrocytes and rapidly (<24h) triggered apoptotic cell death (Fig. 4b). In contrast, ZIKV^u^ only caused an increase in GFAP staining (Fig. 4a, right panel), characteristic of astrogliosis (Yang and Wang, 2015) without changes in cell numbers. Endothelial cells infected with DENV2 or ZIKV^u^(Fig. 4c) displayed disorganized ZO-1 staining, indicating a disruption of tight junctions’ continuity, which is essential to the endothelial barrier role. This disruption may be relevant to the consequences of postnatal infections, including infections in the adult human. Indeed, as shown in Fig. 4d, neuronal cultures could be efficiently infected with ZIKV in a dose-dependent manner. To assess whether virus infection could cause functional impairment, we infected cultures grown on multielectrode arrays. As shown in Fig. 4e, f, infections with DENV2 or ZIKV^u^ induced a rapid loss of electrophysiological activity. Strikingly, DENV2-infected neurons recovered, and resumed activity spontaneously (Fig. 4f), indicating that DENV2 can be neurotoxic, but functional impairment is reversible at the cellular level. In the tested conditions of infection, ZIKV-infected neurons did not recover, eventually degenerating.

**Figure 4.**
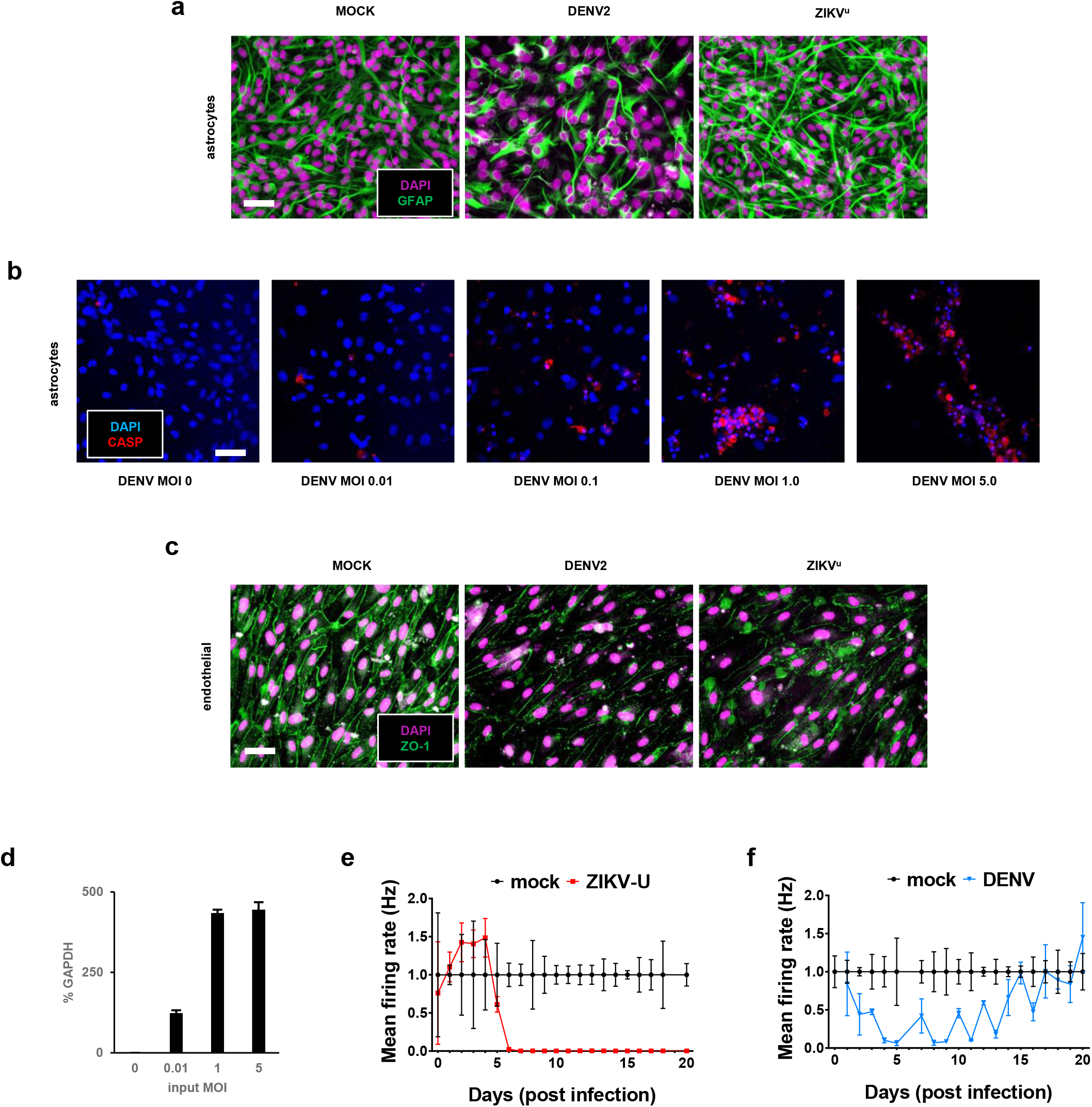
Modeling the postnatal effects of CNS infection. **a.** Astrocytes responded differently to DENV and ZIKV challenges. DENV promoted a loss of viability (middle panel, dapi staining), and an overall decrease of GFAP staining (middle panel, green channel). In contrast, ZIKV did not alter cell numbers (right panel), but triggered a relative increase in GFAP staining intensity, characteristic of astrogliosis; scale bar 40μm. **b**. Astrocytes challenged with increasing amounts of DENV died by caspase-dependent apoptosis (red channel, activated caspase-3 signal); scale bar 40μm. **c**. Another component of the blood-brain-barrier could be modeled with a confluent culture of iPS-derived endothelial cells, forming a network of ZO-1 positive tight junctions (green channel, left panel). Both DENV (middle panel) and ZIKV (right panel) elicited a loss of ZO-1 staining intensity, and disrupted the continuity of the junction network; scale bar 40μm. **d**. A mixed culture of neurons differentiated for 4 weeks could be efficiently infected by ZIKV, in an input MOI-dependent manner (ZIKV RNA as a percentage of *GAPDH);* Values are mean ± s.e.m. (N=2 technical replicates). **e**. A neuronal culture, infected with ZIKV at electrophysiological maturity (6 weeks, day 0 of x-axis), displayed a rapid and irreversible drop in action potential firing rate (red trace) compared to control (black trace). **f**. In contrast, DENV infection resulted in a reversible loss of spontaneous electrical activity (blue trace), reverting to control levels of activity (black trace) after 2 weeks. (e,f) Values are mean ± s.e.m. (N=5 technical replicates for controls, 3 for ZIKV^u^, 2 for DENV)

## DISCUSSION

As is the case with DENV infection, ZIKV has been known to result in a self-limiting and mild disease in most adult patients, with rashes and fevers being the most common symptoms (Petersen et al., 2016). While DENV sometimes leads to a grave hemorrhagic syndrome (Olagnier et al., 2016, Halstead and Cohen, 2015), and in rare cases to neurological damage, maternal ZIKV infection during early stages of pregnancy can often result in fetal neurological abnormalities, ranging from inflammatory scarring with calcifications to severe microcephaly at birth (Coyne and Lazear, 2016, Martines et al., 2016a, Martines et al., 2016b). It is currently unknown how ZIKV crosses all maternal-fetal defenses and barriers to reach the fetal CNS, why virus infection during the 1^st^ trimester is most damaging (Roberts and Frosch, 2016), or why the closely related flavivirus DENV does not cause fetal brain injury. Both viruses are difficult to model in mice, unless interferon defenses are ubiquitously ablated (Sarathy et al., 2015, Miner et al., 2016). Here we show that ZIKV^u^ is under tighter interferon restriction than DENV2 in all studied cells, and that cells which are highly vulnerable, such as neural progenitors, have low baseline defenses and are thus unable to mount an efficient response in the short time before viremia triggers cell death. We also show that microglia can be productively infected and show little cytotoxicity, even when infected with high titers of ZIKV. In contrast, DENV triggers cytotoxic reactions in microglia, as has been reported for other myeloid cells (Martins Sde et al., 2012).

Our results suggest that DENV may very rarely reach the brain, being efficiently restricted by the interferon responses it triggers. In the fetus, immature myeloid cells, and in particular immature microglia, would make poor vectors for DENV as these cells are highly vulnerable to DENV infection. In contrast, ZIKV may have evolved an optimal set of characteristics, allowing it to replicate and propagate in myeloid cells, culminating in infection of microglial precursors without causing serious cytotoxic effects, giving them time to propagate the virus to the tissues they encounter. At the maternal-fetal boundary, cytotrophoblast cells and Hofbauer cells, residing in the chorionic villi, can be infected by ZIKV (Tabata et al., 2016, Quicke et al., 2016). ZIKV can reach these cells through the chorio-allantoic placenta or the highly-vascularized yolk sac splanchnopleur. Our work suggests that microglial precursors migrating between the blood islands of the yolk sac (where they originate) and the embryo proper may represent attractive candidate cells that are susceptible to infection, able to transport the virus to the CNS and mediate infection of neural progenitors (Fig. 3l).

Timing of infection into microglial precursors migrating to the CNS, before closure of the BBB, may be among the mechanisms limiting teratogenicity to the first trimester. *In vivo* evidence of such an event is not currently available, though microglia have been shown to be susceptible to infection in several models, including fetal human slice cultures (Retallack et al., 2016) and non-human primates (Adams Waldorf et al., 2016). The general reliance of mouse models on blunted interferon defenses to observe any pathogenicity of ZIKV hinders direct assessment of the transfer hypothesis. However, striking circumstantial support can be found in a recent study (Vermillion et al., 2017) using immunocompetent mice, bypassing most of the host defenses using intrauterine injections of the virus. In this study, the authors showed that ZIKV, injected at embryonic day 10 (E10), could later be found in the fetal mouse brain, in particular in microglia where it triggered their activation. Remarkably, when injection was performed later at E14, the virus failed to reach the fetal CNS. At E10, yolk sac microglial precursors are entering the CNS, while at E14, this emigration is complete (Ginhoux and Guilliams, 2016). Since microglia, in contrast to NPCs, can still mount a response to the virus, they may conversely help limit viral spread in the CNS parenchyma at later stages, when present in sufficient numbers. We show that DENV2 and ZIKV^u^ may alter the integrity of elements of the BBB, and damage electrophysiologically active neuronal networks, consistent with viral toxicity in the adult (Cao-Lormeau et al., 2016), which may be aggravated in cases of co-infection.

AXL is an attractive candidate receptor for the virus (Hamel et al., 2015, Nowakowski et al., 2016), though AXL ablation was recently found to have no effect on the behavior of ZIKV towards neural progenitors (Wells et al., 2016), and AXL is dispensable for infection in the mouse (Miner et al., 2016). ZIKV may enter cells by redundant use of TAM receptors through apoptotic mimicry (Meertens et al., 2012). The presence of TAM receptors may be particularly relevant in glial cells such as microglia, where the binding of the virus to AXL through the adapter protein GAS6 can downregulate interferon responses (Meertens et al., 2017), allowing the virus to survive encounters with these cells. Other entry mechanisms active in myeloid cells such as microglia include Fc receptors, expressed at high levels, which may allow enhanced endocytosis of virions in the presence of antibodies with partial avidity for ZIKV (Li et al., 2016a, Castanha et al., 2016, Dejnirattisai et al., 2016, Durbin, 2016, Paul et al., 2016). An interesting implication would be that cross-reactive maternal antibodies, some of which may come from previous DENV infections and are present in the fetus, will favor viral infection of primitive myeloid cells such as microglia, further enhancing delivery to the fetal brain (Castanha et al., 2016, Dejnirattisai et al., 2016, Durbin, 2016, Paul et al., 2016). In the adult, prior DENV infections may leave the BBB vulnerable while also enhancing ZIKV infectivity. Ultimately, astrocytes and microglia may become long-term viral reservoirs in the absence of efficient clearing mechanisms, and the consequence of such events remains to be investigated. Our data suggest that type I and II interferon modulation may represent an attractive treatment target, as the teratogenicity of ZIKV may rely on latent vulnerabilities of target cell types (NPCs), evading interferon-dependent eradication in cells such as microglia and astrocytes that may act as Trojan horses or viral reservoirs.

## Acknowledgements

We would like to thank Li-Huei Tsai for providing the iPS-wt5. The authors thank Dongdong Fu, Raaji Alagappan, Tenzin Lungjangwa, and Sarah Elmsaouri, for technical support, and all members of the Jaenisch lab for helpful discussions. Confocal microscopy was performed at the Keck Facility, with the precious help of Wendy Salmon. J.M. received funding from the European Leukodystrophy Association, and a NARSAD Young Investigator Grant from the Brain & Behavior Research Foundation. Y.L. received funding from a Simons Postdoctoral Fellowship, an International Rett Syndrome Foundation (IRSF) Postdoctoral Fellowship, and a NARSAD Young Investigator Grant from the Brain & Behavior Research Foundation. G.B. was supported by H.H.M.I. Work for this project was supported by a grant from the Simons Foundation (SFARI 204106 R.J.), NIH grants HD 045022, R37-CA084198, the ELA foundation, the Emerald foundation, and Biogen (to R.J.) and NIH AI100190 to L.G.

## Author Contributions

J.M., Y.L. and R.J. conceived the project, designed and supervised the experiments, analyzed and interpreted results and wrote the paper with input from L.G. and all other authors. A.O. contributed to all experimental design, performed qPCR assays and immunostaining, and helped write and revise the manuscript. G. B. assisted with organoid section preparation and immunostaining. A.M. and E. R. provided PBMCs, and contributed reagents and expertise on the biology of TAM receptors. A.D. and I.B. prepared viruses, performed Vero infection experiments and contributed to study design and discussion.

## Competing Financial Interests

R.J. is an adviser to Stemgent, a cofounder of Fate Therapeutics and Fulcrum Therapeutics.

## MATERIALS AND METHODS

### Study design

no statistical tests were used to pre-determine sample size. Experimenters were not blinded to the genotypes or culture conditions. All data values were presented as mean +/- SEM. ANOVA analyses were used for comparisons of data with greater than two groups, otherwise a Student t-test was performed. Post-hoc group comparisons were performed with Bonferroni test. A value of p<0.05 was considered significant in all tests. Experiments were performed at least twice.

### Statement of compliance with IRBs

all experiments involving cells from human donors and animals were performed in compliance with established IRB protocols at the Whitehead institute.

### Human ES and iPS cell cultures

Human iPS cell line iPS-wt5 was previously described (REF) and cultured in 5% O2 on mitomycin C-inactivated mouse embryonic fibroblasts (MEFs), in serum-free hESC medium containing DMEM/F12 (Thermo), 20% knockout serum replacement (Thermo), 1% non-essential amino acids (Thermo), 1mM Glutamax (Thermo), 0.1mM β-mercaptoethanol (Sigma) and 12ng/ml bFGF (Thermo). Cultures were passaged with 1mg/ml collagenase type IV (Thermo) every 5-7 days. iPS-wt5 was reprogrammed from a healthy male donor using non-integrative Sendai virus (OSKML). This line is routinely verified for stable expression of Oct3/4, Nanog and TRA1-60, and tested for mycoplasma negativity.

### NGD 0.5X

NGD 0.5X consists of a mixture of 475mL of Neurobasal, 5mL of Gem21-VitA (0.5X, Gemini Bioproducts), 2.5mL of Neuroplex N2 (0.5X, Gemini Bioproducts), 5mL pyruvate 100mM, 5mL Glutamax, 5mL Pen/strep, 5mL NaCl 5M, 1g Albumax I, Biotin 3.5ug, Lactic Acid 85mg, Ascorbic Acid 2.5mg.

### Differentiation to NPCs and maintenance

Differentiation of human ES and iPS cells to neural progenitors in 2-D adherent culture was performed as follows: 4 million human ES or iPS were passaged onto matrigel-coated dishes using PBS w/o Ca^2+^/Mg^2+^, filtered through a 40μm mesh to remove mEFS, and cultured directly in NGD 0.5X medium containing dorsomorphin (2.5μM, Stemgent), bFGF (10ng/mL, Thermo) and Insulin (additional 10ng/mL) until super-confluent. bFGF and Insulin were removed after a week, and NGD 0.5X + dorsomorphin was replaced every day for 10 days. Cells were subsequently passaged 1:1 with PBS^-/-^ when rosette lawns were observed throughout the culture. Rho-associated protein kinase (ROCK) inhibitor Y27632 (10mM, Stemgent) was added to the medium during each of the first 3 passages. Initial passaging at no more than 1:2 ratio, followed by 1:3 to 1:6 every 5 days. After the intial passage, neural progenitors were expanded and maintained in NGD 0.5X with 10ng/ml bFGF, and addition of 10ng/mL human Insulin (NGM).

### Differentiation into neurons and astrocytes

NPCs were differentiated into neurons and glia by removing FGF and culturing in NGD 0.5X without retinoic acid addition (NGD contains no additional retinoid). To initiate differentiation, NPCs were plated in a 35mm dish on 1% matrigel at 5×10^5^/cm^2^ and fed 5mL of NGD every 2–3 days. At 4 weeks, neurons appeared in the culture and a final dissociation was performed. The culture was dissociated in the presence of 0.05% DNase I by incubation with Accutase (Stem Cell Technologies) for 30’ at 37°C with gentle agitation (bacterial rotator), re-suspended in chilled HBSS/0.1% BSA and filtered through a 40μM mesh before being centrifuged at 100g for 10’ through a cushion of 4% BSA to remove cellular debris. For 2D cultures cells were re-plated at 2×10^5^/cm^2^ on PDL-matrigel coated plastic or glass. Spheroid formation was initiated by re-aggregating 3×10^4^ NPCs per well in 96-well ultra-low attachment plate (corning), allowed to differentiate in suspension. PSA-NCAM-and A2B5+ glial progenitors were derived from multipotent NPCs, and cultured in NGD 0.5X medium with the addition of EGF (20ng/mL) and bFGF (20ng/mL) as immature astrocytes. Further maturation was achieved with the withdrawal of FGF and EGF, and addition of 5% FBS or CNTF (10ng/mL).

### Differentiation into microglia-like cells (based on Muffat et al., 2016b)

Colonies were treated with collagenase IV (1.5mg/mL) and mildly triturated to form a suspension of uniform clumps, transferred directly to 5mL NGD 0.5X + 10ng/mL IL-34 + 20ng/mL CSF1, in ultra-low attachment 6-well plates (corning). 6 confluent wells (~3×10^6^ cells) were pooled into one suspension well. Embryoid bodies were monitored for appearance of compact phase-bright neuralized spheroids, as well as expanding cystic yolk sac embryoid bodies. Every 5 days, EBs were gently triturated to shear off loose cells of interest, settled, and the supernatant placed in a single well of a Primaria 6-well plate, left to adhere for an hour. Unattached cells and small EBs were washed with DMEM/F12+0.1% BSA, and wells were fed with fresh medium. Attached cells were monitored for morphological characteristics of microglia/microglial precursors (compact nucleus, vacuoles, membrane ruffles, motility), and wells from 6 consecutive productions (one month) were pooled to constitute one mixed birthdate population. Infections were performed in NGD 0.5X + 10ng/mL IL-34 + 10ng/mL CSF1.

### Organoid infection assay

Neural progenitors were generated as described. After a month of generation and maintenance (corresponding to passage 3-5), 3×10^4 cells were re-aggregated in 96-well ultra-low adherence wells to generate neural spheroids. These organoids were left to grow and differentiate in NGD 0.5X for 7 days. After a week, organoids kept in 96-well format were infected with ZIKV (MOI 1 matching the surface cell number, assuming 15μm diameter per cell), and observed for 7 more days in medium without growth factors, measured for size variation. Alternatively, microglia were infected at an MOI of 1, and 10^3 infected cells were added 24 hours after infection to a single organoid. Organoids were collected at various time point and RNA was prepared. After 7 Days, organoids were fixed, cryo-sectioned, and stained for ZIKV and IBA1.

### Primary human monocytes

PBMCs were obtained from healthy donors and purified by differential centrifugation, then cultured on Primaria in the medium used for microglial culture. Human tissue used for the isolation of primary cells is derived from donors who have signed informed consent by the donor themselves.

### In situ cell counting

Cells were imaged in phase contrast. FiJi Gaussian blur was applied to the images, followed by background subtraction and thresholding. The particle counting function was used to count the number of cells remaining in the well after 72 hours, post exposure to increasing MOI of initial inoculum.

### Electrophysiology

Neural progenitors were differentiated in NGD 0.5X for 4 weeks, and plated on PEI coated 12-well multielectrode arrays (axion biosystems) to be analyzed on the Maestro recording system. Recording was performed daily for 5 minutes. The neural signal processing routine was used to generate timestamps for each field potential recorded at each electrode. Data is presented as well-based mean firing rate, normalized to control wells (display as 100%).

### Imaging

Cells and tissues were fixed with ice-cold methanol or 4% paraformaldehyde in PBS. After PFA fixation, permeabilization was performed with PBS containing 0.3% triton. Fixed and permeabilized cells were blocked with 3% normal donkey serum. Primary antibodies were against Flavivirus (Santa Cruz), GFAP (Abcam), Sox2 (R&D Systems), PU.1 (cell signaling, 1:500) and Iba1 (abcam, 1:500) and visualized by *ad hoc* secondary antibodies conjugated with Alexa 488, 568, 594, 647 (Life Technologies, 1:1000), followed by counter-staining with DAPI. Phase contrast images of infected cultures were performed on an EVOS microscope. Fluorescent confocal images were acquired on a Zeiss LSM 700 at 20X magnification.

### RNA extraction, reverse transcription and quantitative PCR

Cells were homogenized and total RNA extracted using the RNeasy Micro or Mini kit (Qiagen) following the manufacturer’s instructions, and vRNA was extracted from conditioned medium using the QIAmp viral RNA kit. Total RNA concentrations were measured using NanoDrop ND-1000 spectrophotometer. RNA was reverse transcribed into cDNA using Superscript III reverse transcriptase (Invitrogen) with random hexamer primers. Transcript abundance was determined by quantitative PCR using SYBR Green PCR mix (Applied Biosystems) or Taqman assay.

### Cytokine profiler

5×10^4 microglia were plated, and infected after 24 hours at MOI 1. Inoculum was left in contact with the cells for 12 hours, washed off, and 1mL of fresh medium was allowed to condition from 12 to 24 hours post infection. 400μL of this supernatant were added to a prepared cytokine antibody panel membrane, per the manufacturer’s instructions (RnD Systems, kits ARY005). After incubation, the membranes were revealed with ECL reagent.

### Interferon modulation

20ng/mL IFN-γ or 100ng/mL B18R were added 2 hours before viral infection, and 24 hours after cell plating. The inoculum was left in contact with the cells for 12 hours before being washed and replaced with fresh corresponding media.

### Viral stocks and infection protocol

ZIKV strain MR766 (Uganda) was obtained from ATCC, and expanded in Vero cells or C6/36 mosquito cells. Dengue virus serotype 2 strain 16681 was expanded in C6/36 mosquito cells. To establish titered viral stocks, virus-containing supernatant was harvested and viral titer was determined by infecting Vero cells, followed by quantitative flow cytometry analysis of Flavivirus envelope immuno-staining to calculate Vero cell infectious units (Lambeth et al., 2005). MOI used to infect 2D and 3D culture was calculated based on Vero cell infectious units. For direct infection of organoids, the number of cells on the surface was estimated from the diameter of the 3D organoid and was used to calculate the amount of virus applied. Viral inoculum was washed off and replaced after 24hours with fresh medium and size was recorded at day 0 and day 7 (masked area of projection).

### >Viral transfer

For Vero cells reinfection (Fig. 3d), 1mL supernatant from an equivalent number of cells of various types, infected at MOI1 72 hours prior, was added to a confluent lawn of Vero cells in 24-well format. After 24hours cells were dissociated, fixed and analyzed by FACS for viral envelope positivity. For re-infection of NPCs by microglial supernatant (Fig. 3e), 100uL of 72hpi supernatant from 5×10^4 microglial were added to 1mL of medium covering 5×10^5 NPCs. Supernatant was replaced 24 hours later and cells were stained in situ with an antibody against viral envelope proteins and counterstained with DAPI.

